# Robustness of spike deconvolution for calcium imaging of neural spiking

**DOI:** 10.1101/156786

**Authors:** Marius Pachitariu, Carsen Stringer, Kenneth D. Harris

## Abstract

Calcium imaging is a powerful method to record the activity of neural populations, but inferring spike times from calcium signals is a challenging problem. We compared multiple approaches using multiple datasets with ground truth electrophysiology, and found that simple non-negative deconvolution (NND) outperformed all other algorithms. We introduce a novel benchmark applicable to recordings without electrophysiological ground truth, based on the correlation of responses to two stimulus repeats, and used this to show that unconstrained NND also outperformed the other algorithms when run on “zoomed out” datasets of ~10,000 cell recordings. Finally, we show that NND-based methods match the performance of a supervised method based on convolutional neural networks, while avoiding some of the biases of such methods, and at much faster running times. We therefore recommend that spikes be inferred from calcium traces using simple NND, due to its simplicity, efficiency and accuracy.

## Introduction

Two-photon calcium imaging can be used to monitor the activity of populations of up to 10,000 neurons^1^. Nevertheless, calcium-sensitive fluorescence signals are an indirect readout of cellular activity. Therefore, accurate and well-calibrated data processing methods will be required to make optimal use of this activity^2–10^. One important problem is developing methods for spike detection: inferring the times of action potentials from the fluorescence traces. The earliest such methods rely on spike deconvolution algorithms, which infer a spike train under the assumption that the fluorescence trace represents an approximate convolution of the underlying spike train with the cell’s calcium response^10^. This is often a good approximation^11^, though situations exist when it breaks down. More complex spike deconvolution algorithms take these extreme cases into account^3^.

Recently, a new approach to spike detection, based on supervised learning, has been claimed to outperform several existing deconvolution algorithms^9^. Supervised algorithms learn to solve the spike detection problem by training on “ground truth” data where spike times are also measured electrophysiologically. In principle, such methods should give the most accurate results: however, they may generalize poorly to “out-of-sample” data, i.e. recordings made under different conditions to the available training data.

Since the supervised approach was first introduced, ground truth data released as a public benchmark (“spikefinder”) has allowed the comparison of several old and new algorithms. All three winning algorithms (called “Elephant”, “Purgatorio”, “convi6”) employ supervised methods based on convolutional neural networks, and appear to outperform all NND-based methods (“oopsi”, “Suite2p”, “MLspike”). However, this performance may be due to specific design features of the spikefinder challenge, rather than true improvements in spike deconvolution quality. First, the spikefinder benchmarks are run on in-sample data, and may thus not reflect generalization performance to new recordings. Second, multiple metrics for the similarity of decoded and actual spike trains are possible; because supervised methods can be trained to optimize the particular benchmark used, they will have an advantage over unsupervised methods, unless the latter are also optimized for the particular quality metric used.

We show here that non-negative deconvolution (NND) – with very simple parameter settings, and using the fast OASIS implementation^4^ – outperforms supervised algorithms, when it is: 1) evaluated on out-of-sample data, and 2) adapted to the performance metric of the spikefinder challenge. In addition, we find that NND is highly robust to assumptions on the assumed shape of the calcium response to single spikes (henceforth called a "kernel"), such that a simple decaying exponential kernel performs better than more biologically accurate kernels that include a rising time segment, and even performs better than kernels estimated directly from ground truth data. Moreover, large changes to the timescale of the exponential kernels did not affect performance significantly, and optimizing these timescales for each cell actually hurts performance. Finally, we propose a new benchmark that can be used without electrophysiological ground truth, and show that simple NND again outperforms other algorithms on this benchmark.

## Results

For the first set of benchmarks, we considered two main classes of datasets with simultaneously-recorded ground truth electrophysiology, which we will refer to in short as “GENIE” and “SPIKEFINDER”. The GENIE collection consists of five datasets recorded by the GENIE project^11–14^, that have been used in the original descriptions of the GCaMP calcium sensors and their red variants and are available on CRCNS.org^14^. The SPIKEFINDER collection consists of the five datasets analyzed in Theis et al, 2016, made publically available as part of the “spikefinder” challenge (spikefinder.codeneuro.org). Surprisingly, the two state-of-the-art algorithms, “oopsi” and “stm”^9,10^, have different performance on the two GT datasets, with oopsi winning on GENIE and stm winning on SPIKEFINDER.

To determine deconvolution performance, we need to establish a set of comparison metrics between the ground truth spike trains and each algorithm’s output. Standard metrics compute the similarity of the true and inferred spike trains after binning both in a preset window size, which is typically small (i.e. 40ms in the spikefinder challenge). After binning, one typically computes the correlation between the true and inferred spike train. Such metrics may not be well suited for spike trains, which are very sparse quantities. For example, if an inferred spike is offset by just one bin from a GT spike, it will be counted as a complete miss by the correlation metric, similar to temporal mismatches of several bins. Supervised algorithms, unlike unsupervised ones, have automatic protection from this effect, as they are directly trained to minimize the correlation metric. Unsupervised algorithms can be adapted to perform well by the correlation metric by smoothing their output temporally, which reduces the effect of temporal mismatches between true spikes and deconvolved spikes. In addition, a temporal offset might be required for some datasets (for example due to hardware synchronization issues), which can be learnt automatically by the supervised methods, but has to be inferred post-hoc in the unsupervised algorithms.

A short segment of fluorescence from a cell recorded with ground truth is shown in Figure 1a, together with the reconstructions obtained with three unsupervised models (the quantity ***s ∗ k*** in equation 1). All three models track the large fluorescence changes, but some of the smallest changes are only tracked by the unconstrained model (NND), which is relatively less constrained than the L0- or L1- penalized methods. For example, the L1-penalized model overly penalizes single spike events in some cases, reducing their amplitude relative to other single or multi spike events (Figure 1b). Nonetheless, for this cell, all three deconvolution methods returned roughly similar spike trains, which correlated well with the known ground truth electrophysiology (Figure 1c).

**Figure 1.**
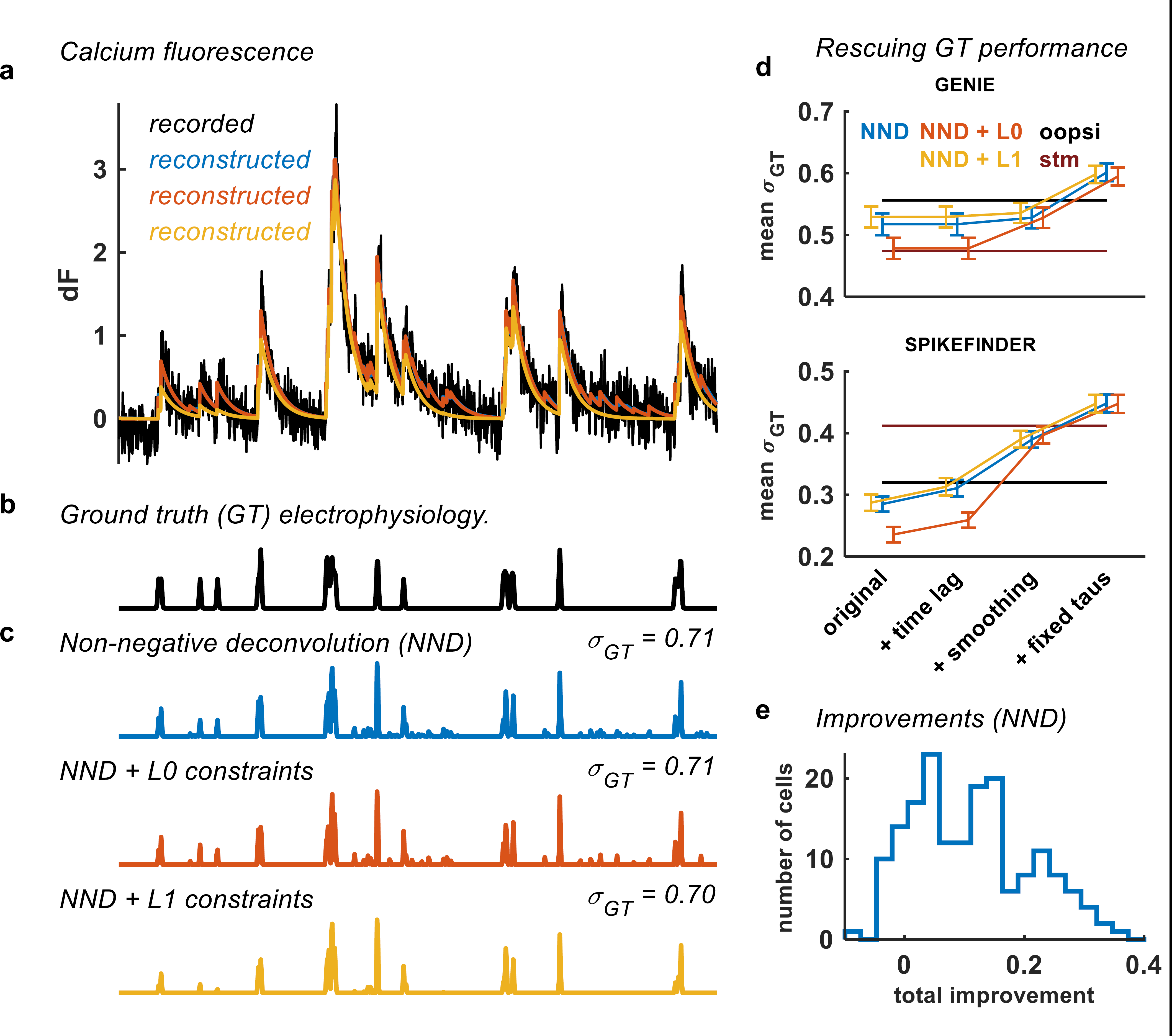
Performance with ground-truth electrophysiology. (a) Example fluorescence recording of a neuron recorded by the GENIE project. The model trace reconstructions are shown in color (blue is NN, red is NN + L0, yellow is NN + L1).
(b) Simultaneous ground truth electrophysiology for the neuron shown in **a**.
(c) Deconvolved traces using three NND models.
(d) Correlation *σ_GT_* between deconvolved and ground truth spike trains, with various processing stages included, averaged over cells separately for datasets from the GENIE project *(Chen et al., 2013)* and for SPIKEFINDER datasets *(Theis et al., 2016).* The average values for stm and oopsi are taken from the spikefinder challenge (http://spikefinder.codeneuro.org).
(e) Distribution across cells of the total improvement provided by the post-processing of outputs from unconstrained NND.

### Simple non-negative deconvolution outperforms the state-of-the-art results

There has recently been a proliferation of calcium deconvolution algorithms^2–10^ some of which have provided their code publicly. However, little effort has been made to compare the performance of these algorithms to each other, with the notable exception of Ref. ^9^, who concluded that supervised algorithms, trained on available ground truth data, perform better than the more routinely used unsupervised algorithms.

We investigated this claim on the same datasets used by Ref. ^9^, which were since made available publicly in the “spikefinder” challenge. We indeed found that the supervised algorithms performed better than the L0 and L1-penalized algorithms, when evaluated by the correlation metric (Figure 2c, “original”). However, we found that unsupervised algorithms became superior after very simple modifications.

**Figure 2.**
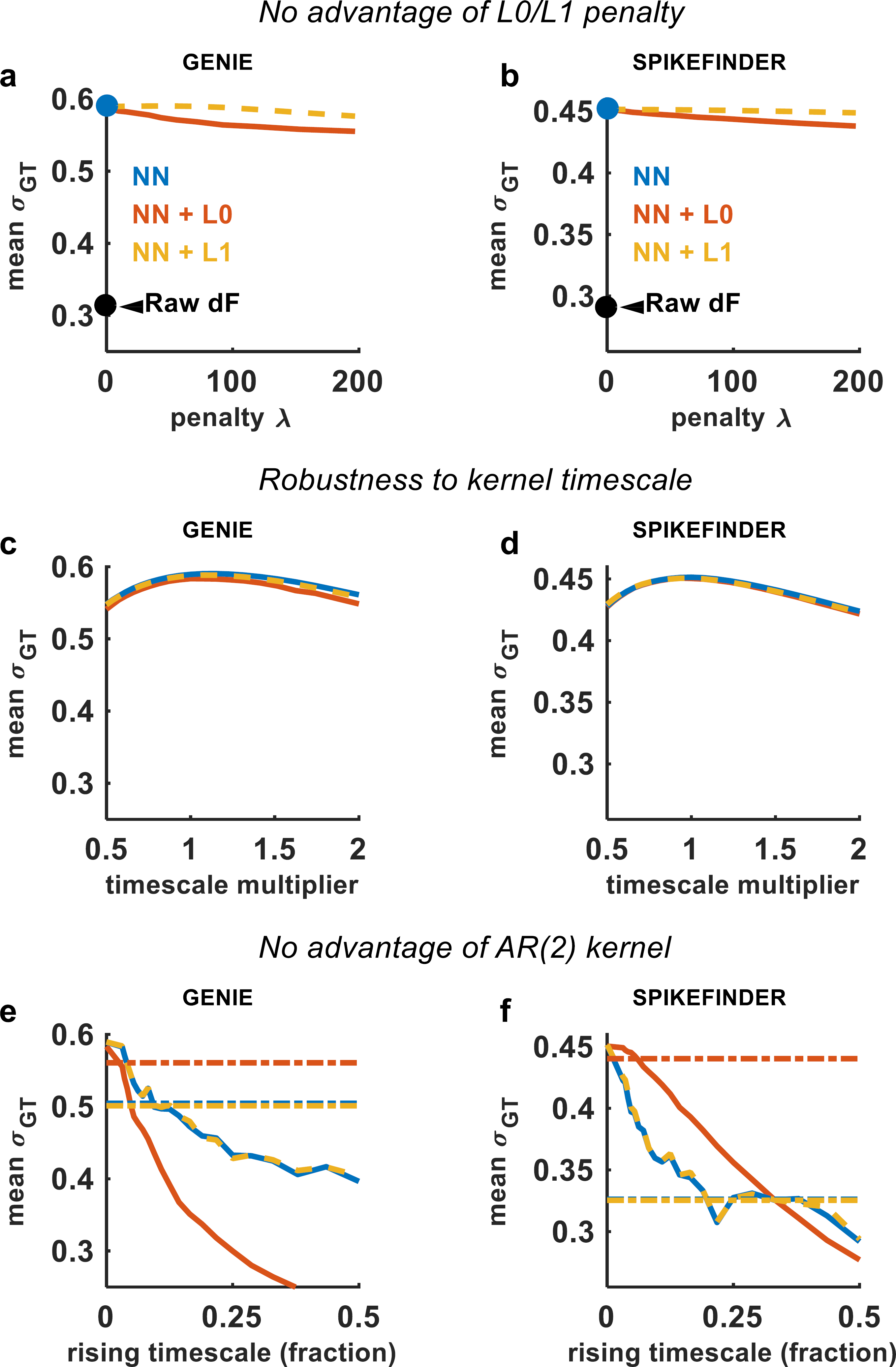
Robustness to changes in parameters. Effect of parameter values on mean correlation *σ_GT_* between deconvolved spike trains and ground truth electrophysiology. (ab) The penalty on sparsity was varied, for both L0 and L1-penalized models. The optimal value of 0 corresponds to unconstrained NND.
(cd) The kernel timescales were varied; this had little effect but did not significantly improve performance over values in the literature
(ef) A second kernel timescale was introduced and varied, to model the rise time of the fluorescence following a spike. The kernels were defined as a difference of exponentials, which defines a subclass of AR(2) kernels. The AR(2) version of OASIS was used for the L1-penalized method. Performance with ground truth-derived kernels is also shown as dotted lines (see Methods). The optimal rising timescale was 0, corresponding to a simple exponential kernel.

First, we chose an appropriate timelag parameter for a subset of datasets where the timing of the ground truth spikes was not perfectly synchronized with the fluorescence (Figure 1c, “+time lag”). This parameter was chosen to maximize the correlation with the ground truth spikes, for each of the ten datasets separately. The inferred timelag was 0 for all GENIE datasets, and ranged between -2 and 3 for SPIKEFINDER datasets.

Second, we applied smoothing to the deconvolved traces, which reduced the effect of the correlation metric (Figure 1c, “+smoothing”). The smoothing was performed with a Gaussian-shaped kernel with a standard deviation of two samples for all GENIE datasets, and 8 samples for all SPIKEFINDER datasets. These values were empirically found to perform well. Note that all datasets have been upsampled at 100Hz, and are benchmarked at 25Hz.

Finally, we did not allow the algorithms to estimate the best fit calcium kernels, or the kernel’s decay timescale, as we found that all methods failed to recover appropriate parameters. Instead, we fixed the timescales of the calcium kernel to be approximately the measured values from the literature^11–13^ (Figure 1c, “+fixed taus”). For simplicity, we divided all sensors into a fast, a medium and a slow category, and used timescales of 0.5, 1 and 2 seconds for the three categories. As we show below, the precise values for these timescales were not critical.

These improvements, together, increased the benchmark performance for nearly all cells (Figure 1d), and surpassed both the supervised and unsupervised “state-of-the-art” approaches submitted on the website spikefinder by their developers (stm and oopsi). Furthermore, the best performing model in the benchmark was unconstrained NND, with the L0- and L1-based methods slightly lagging behind (Figure 1d).

### Robustness of non-negative deconvolution

These results suggest similar performance levels from different regularization methods for unsupervised deconvolution. We tested the relation between the algorithms more systematically, by varying the regularization parameter *λ*. When *λ =* 0, both the L0 and L1-based algorithms are equivalent to the unconstrained approach. We found that for both algorithms, performance was best when *λ =* 0, corresponding to unconstrained NND (Figure 2ab).

All three unsupervised algorithms were robust to large changes in the shapes of the calcium kernels. Lengthening or shortening the assumed timescale of the calcium indicators by a factor of 2 did not significantly affect performance (Figure 2cd). Furthermore, adding another component to the kernel (a “rising” timescale) did not improve performance (Figure 2ef). Finally, performance was not improved by using the “ground truth” kernel, obtained directly by regressing the fluorescence onto the ground truth spikes (Figure 2ef, dotted lines).

### Ground truth -free benchmarks using stimulus responses

The GENIE and SPIKEFINDER datasets contain several tens of cells each. However, the conditions in which these cells were recorded might be different from conditions in many experiments. To assess how spike deconvolution methods perform in such circumstances, we developed a novel benchmark that does not require ground truth electrophysiology (Figure 3a).

**Figure 3.**
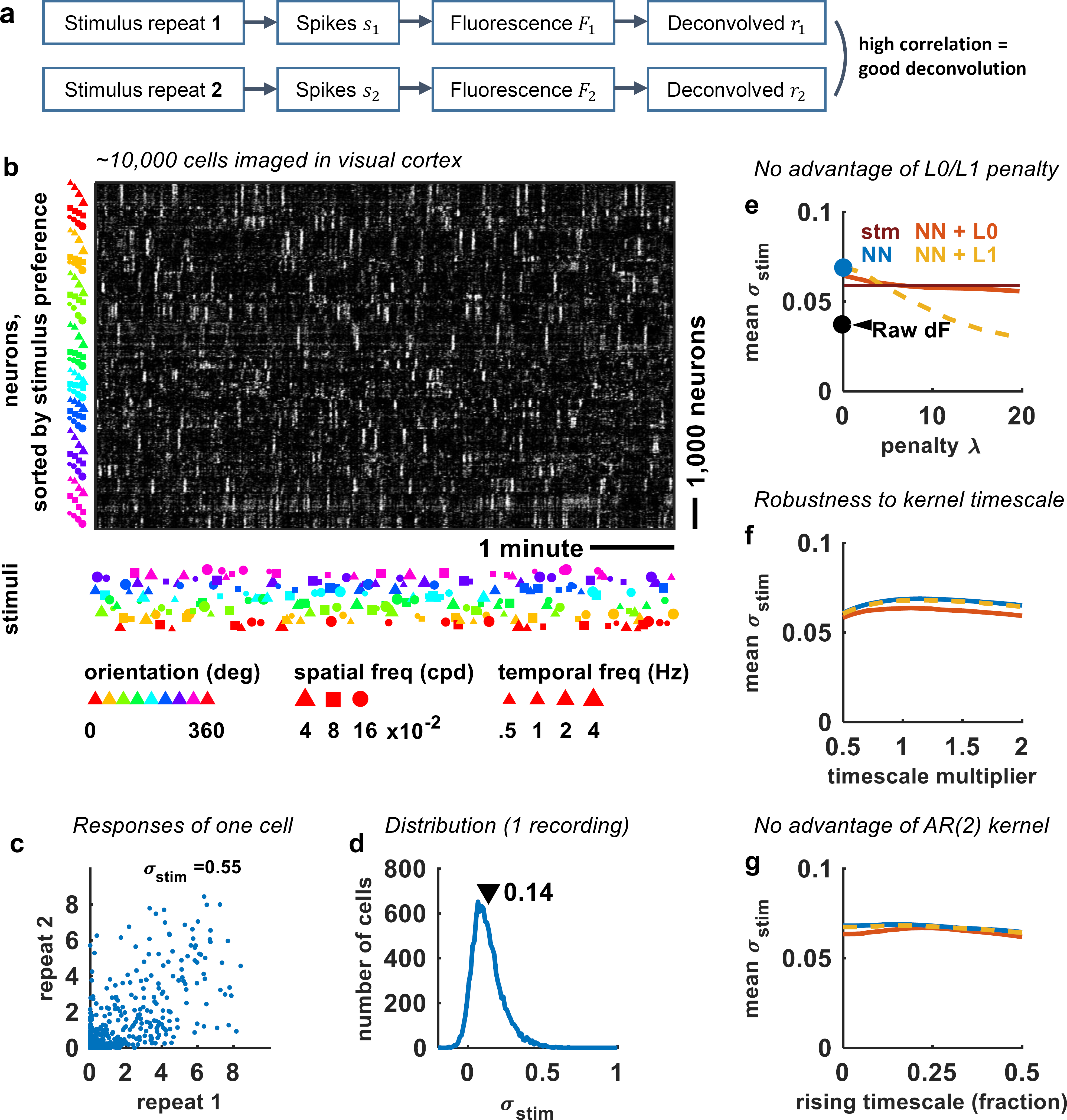
Benchmarking without ground truth electrophysiology. (a) Example deconvolved neural responses (NND algorithm) from ~10,000 simultaneously recorded neurons, sorted by their preferred stimulus. The stimuli shown were drifting gratings with one of 8 directions, one of 3 spatial frequencies and one of four temporal frequencies.
(b) Correlation between neural responses to half of the stimulus presentations, versus the other half of the stimulus presentations.
(c) Distribution of correlation coefficients (Spearman; equivalently signal variance fraction, see Methods). The mean of these coefficients is used in **d-f** as a benchmark of deconvolution performance (higher is better).
(d) Deconvolution performance as the penalty on sparseness was increased, for L0 and L1-penalized models, as well as for raw non-deconvolved data, and for the stm model from Theis et al, 2016.
(e) Deconvolution performance as the timescales are increased or decreased by a fractional amount.
(f) Performance with AR(2) kernels that include a non-instantaneous rise with varying timescales.

**Our new benchmark is to maximize the Spearman correlation *σ_stim_* of deconvolved responses to** two repeats of an ensemble of stimuli: *σ_stim_* = corr*(Ŝ_repeat1_, Ŝ_repeat2_)*, where *Ŝ_repeat1_* and *Ŝrepeat2* are the binned responses to *N* stimuli. The *N* stimuli cannot be identical, and must be presented in randomized order on each of the two repeats. It can be shown that *σ_stim_* approximates the proportion of signal-related variance contained in the trial-averaged responses to this stimulus ensemble, with 1*-σ_stim_* representing the proportion of noise variance (reflecting a sum of biological and measurement noise; see Methods). Since the signal can only originate in the true spiking *s, σ_stim_* captures the ability of the deconvolution to reconstruct the true spiking. Deconvolution can fail to capture the signal variance in *s* in one of two ways: 1) failing to distinguish spikes from noise, or 2) failing to temporally localize spikes to the correct stimulus bins.

For this analysis, we used some of our own datasets, in which the responses to an ensemble of full field drifting grating stimuli were simultaneously-recorded in ~ 10,000 cells from primary visual cortex of awake mice. A raster plot of these responses (deconvolved by NND) is shown in Figure 3b. Responses of single cells were noisy (Figure 3c), but most cells had positive Spearman correlations *σ_stim_* between the two stimulus repeats (Figure 3d). Taking *σ_stim_* as a measure of deconvolution performance (higher is better), we repeated the types of analyses from Figure 2, for 6 datasets recorded from 4 mice. Again we find that NND without constraints performs better than the constrained versions, and again outperforms the supervised algorithm introduced by Theis et al, 2016 (Figure 3e). We also find that the unsupervised deconvolution methods are robust to the kernel timescale up to a factor of 2 (Figure 3f), and that the AR(2) kernel does not help performance (Figure 3g). We conclude that this new benchmark reinforces the results obtained on data with simultaneous electrophysiology. In addition, we note that this method can be easily applied to other recordings, allowing users to benchmark multiple approaches on their own data.

### Comparisons with convolutional neural networks

We have shown that unsupervised methods, when adapted to the correlation benchmark, outperform the supervised approach described in Theis et al., 2016, on the same ground truth data used there. However, it is possible that more advanced supervised methods perform better still. In the recent “spikefinder” challenge, convolutional neural networks (CNNs) appear to outperform all other methods. However, to successfully apply CNNs on standard recordings without ground truth, it must be shown that CNNs generalize to datasets not included in the training set. We thought this generalization might be imperfect, because CNNs, like all supervised methods, can overfit to the specific statistics of the ground truth datasets.

We evaluated the generalization performance of CNNs, by training the “Elephant” method on nine of the ten available datasets, and testing it on the tenth. We found that under this testing protocol, the performance of supervised CNNs and unsupervised NND was nearly identical, both on the GENIE datasets (CNN: 0.44, NND: 0.45) and on the SPIKEFINDER datasets (CNN: 0.59, NND: 0.60), although there were slight variations in performance from neuron to neuron (Figure 4a). We also found very similar performance in the ground-truth free benchmarks we just introduced above (CNN: 0.067 vs NND: 0.071), with some variation from neuron to neuron (Figure 4b).

**Figure 4.**
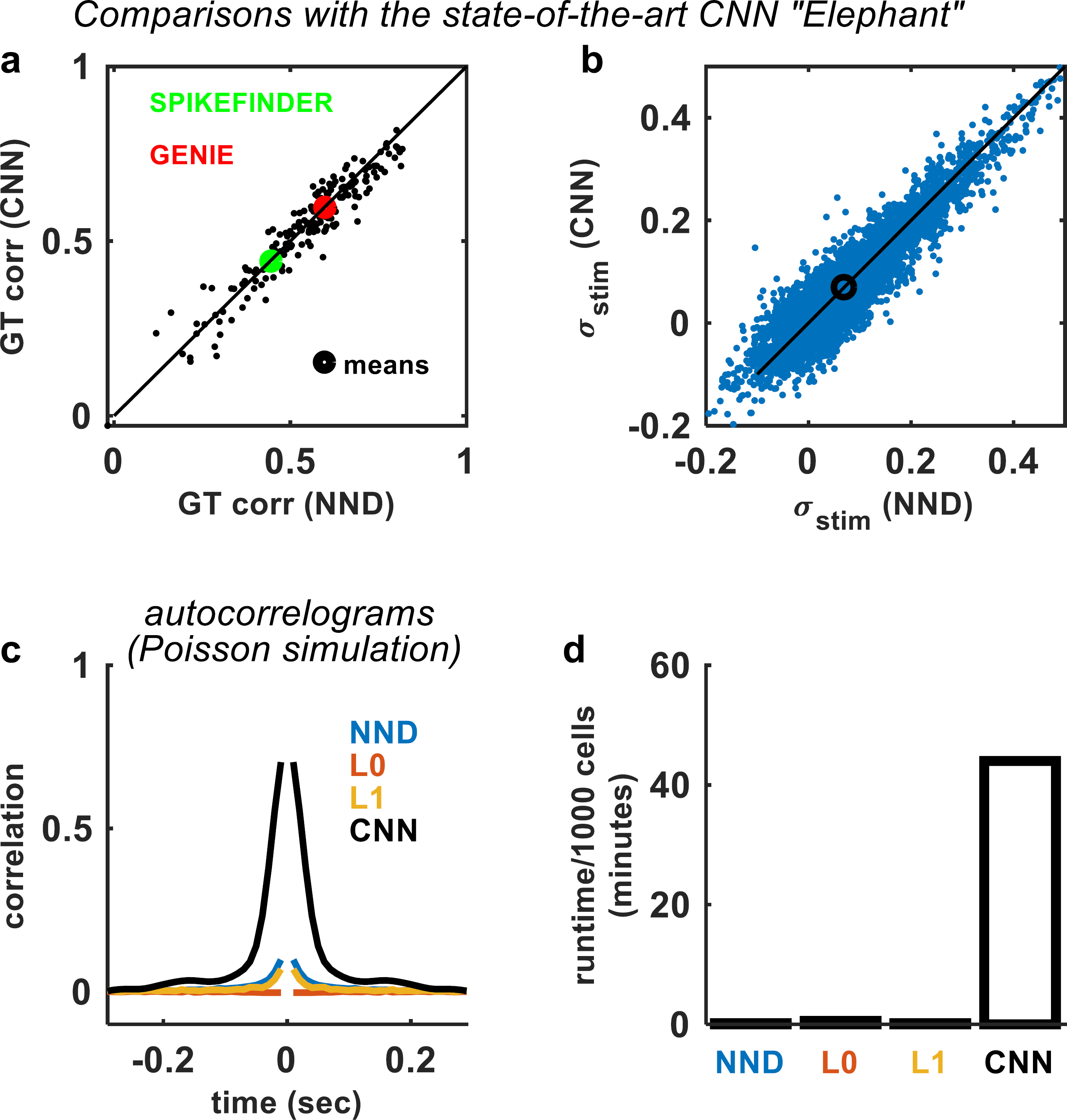
Non-negative deconvolution matches the performance of supervised, convolutional neural networks. (a) Spike detection performance for all cells with ground truth electrophysiology. The means for all GENIE and all Theis et al, 2015 datasets are shown as circles. Each CNN was trained on all but one of the 10 datasets, and tested on the remaining dataset. Compare with Figures 2,3.
(b) Signal variance of CNN trained on all 10 datasets, and tested on our data using the repeat similarity benchmark (Figure 3).
(c) Auto-correlograms of deconvolved spike trains from simulations with Poisson ground truth statistics. The CNN approach heavily biases the statistics of the inferred spike trains. The L0 and L1 method were run with same parameters as in Figure 2 (*λ* = 10 and 100 respectively).
(d) Runtimes of CNN on a high-end GPU (GTX 1080), compared with runtimes for OASIS (NND and L1) and the L0 method on a standard CPU (Core i7).

The similarity in performance might appear at odds with the results of the spikefinder challenge, where the CNN methods outperformed unsupervised approaches by ~10% (CNN: ~0.46 vs NND+L0: ~0.43). However, for that challenge the CNN was tested on within-class data, thus being able to take advantage of the particular statistics of spiking and fluorescence for each recording. One such statistic is the autocorrelogram structure of the spike trains, which was far from Poisson, reflected either stimuli were presented during some of the recordings, or the structure of spontaneous activity in the recorded neurons. The autocorrelogram structure can be used by supervised approaches to perform better spike prediction. However, this strategy is undesirable, because it will enforce the properties of the training data on new data, potentially leading to an erroneous scientific conclusion that all recorded neurons share the same temporal dynamics as the neurons used to train the algorithm.

To demonstrate this transfer of constraints between training and test data, we simulated spike trains with Poisson statistics (flat auto-correlograms), and generated fluorescence traces from them with a calcium decay timescale of 1 second. The deconvolved spike trains using CNNs had a large, spurious auto-correlation at short timelags, which was much less pronounced in methods based on OASIS (NND and NND+L1) and absent using the L0-based method (Figure 4c). The “black-box” nature of CNN algorithms raises a further concern that other features of the training data may be erroneously imposed on new data, in ways that are unknown to the user.

Finally, another disadvantage of complex CNNs is speed: even using a high-performance GPU (GTX 1080), the method is two orders of magnitude slower than all unsupervised methods we tested, which can run efficiently on standard CPUs (Figure 4d).

## Discussion

We conclude that the performance of simple NND-based deconvolution algorithms matches or exceeds all tested alternatives, and that L0/L1 penalties provided no advantage. NND was robust to changes in kernel timescale or shape, with values taken from the literature providing close to optimal performance. Automated identification of kernel parameters appeared to be counterproductive, resulting in significantly mismatched parameters that impaired performance. While supervised methods gave apparently superior performance in previously reported benchmarks, this reflected their ability to optimize particular evaluation metrics, and compensate for phenomena such as synchronization lags within single example datasets; when tested out-of-sample, against unsupervised methods with appropriate compensatory mechanisms, we found their performance to be inferior. We therefore recommend unconstrained NND, with fixed calcium decay timescale.

OASIS provides a very efficient algorithm for performing this deconvolution^4^. In Suite2p, the calcium processing pipeline that we maintain, we provide wrappers for the OASIS toolbox, and additionally include the L0-based deconvolution code, which may provide advantages for some cases, such as avoidance of auto-correlation bias (Figure 4c). It remains to be seen if new methods can significantly outperform simple NND. However, marginal improvements of future approaches might be overshadowed by the simplicity and interpretability of the NND approach.

## Materials and methods

### Imaging in visual cortex

All experimental procedures were conducted according to the UK Animals Scientific Procedures Act (1986). Experiments were performed at University College London under personal and project licenses released by the Home Office following appropriate ethics review.

The experimental methods were similar to those described elsewhere^15^. Briefly, surgeries were performed in adult mice (P35–P125) in a stereotaxic frame and under isoflurane anesthesia (5% for induction, 0.5-1% during the surgery). During the surgery we implanted a head-plate for later head-fixation, made a craniotomy with a cranial window implant for optical access, and, on relevant experiments, performed injections of the GCaMP6m virus with a beveled micropipette using a Nanoject II injector (Drummond Scientific Company, Broomall, PA 1) attached to a stereotaxic micromanipulator. Viruses were acquired from University of Pennsylvania Viral Vector Core. Injections of 50-200 nl virus (1-3 × 10^12^ GC/ml) were targeted to monocular V1, 2.1-3.3 mm laterally and 3.5-4.0mm posteriorly from Bregma and at a depth of L2/3 (200-400 μm). Some mice were transgenic and expressed tdtomato in certain cell classes. However, we did not use that information here.

### NND Model

NND models infer the most likely spike train **s**(*t*), given the fluorescence timecourse **F**(*t*), and a response kernel **k**. Models based on deconvolution define a cost function of the form

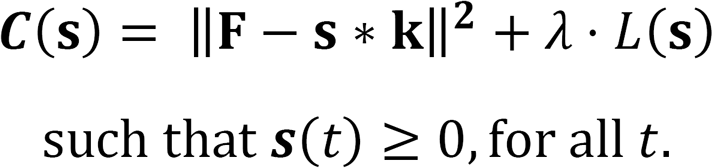

Here, **s ∗ k** describes a temporal convolution of a positive timecourse ***s*** and the kernel ***k***, and *L*(**s**) describes a penalty function on the inferred spike trains.

We tested a suite of three unsupervised spike detection methods. The first is an approximate optimization algorithm where *L*(**s**) = ‖**s**‖_0_ is the L0 norm, i.e. the number of non-zero entries in **s** (code available at github.com/cortex-lab/Suite2P). The L0 penalty enforces the constraint that the inferred spike trains should be very sparse, because neurons fire rarely. The second method has *L*(**s**) = ‖**s**‖_1_, the L1 norm, and chooses the kernel from a parametrized class of functions^4^. The third unsupervised model is unconstrained non-negative deconvolution (NND), with *L*(**s**) = 0. For our initial analysis, we chose the sparsity penalty *λ* for both the L0 and L1 methods in such a way as to output spike trains with similar levels of sparsity: only ~5% of the deconvolved samples were non-zero, for data sampled at 100Hz. We then varied the sparsity penalties, as well as several other parameters, to understand how they influence performance. We compared performance of these algorithms against two supervised methods: the method of Theis et al^9^, and also a publicly available convolutional neural network algorithm (code from https://github.com/PTRRupprecht/Spikefinder-Elephant/tree/master/elephant).

### L0-based spike deconvolution

We obtained a fast deconvolution algorithm by developing a novel optimization procedure for a standard spike generation model. Our optimization is an extension of a well-known algorithm called “matching pursuit”^16^. Briefly, the matching pursuit algorithm identifies putative spikes by their similarity to the calcium kernel. It then subtracts the kernel scaled with an appropriate factor (the “spike amplitude”) from the location of the identified spikes. On successive iterations, more putative spike locations are identified greedily, and their activity subtracted off. This continues until no new spikes can be introduced, because they do not have large enough amplitudes to explain a significant portion of the variance.

This basic matching pursuit algorithm is limited by local minima, because the greedy procedure cannot always resolve nearby spikes that have overlapping calcium activity. To mitigate this problem, we add an extra step at each iteration, where we allow existing spikes to change their location and/or magnitude to better account for the calcium trace. The optimal changes can be calculated exactly and efficiently, allowing “old” spikes to adjust their locations and activity in the context of “new” spikes, thus reaching more accurate solutions. We note that another algorithm for solving a similar problem has recently been proposed^5^. Although this algorithm does not impose positivity, and is slower than our approach, it obtains an exact solution for their respective problem.

### Stimulus-related variance from two repeats

Here we show that the correlation *σ_stim_* of a neuron’s responses to two stimulus repeats is equal to an unbiased estimate of the signal variance of that neuron, as a fraction of its total variance. Define a neuron *n*’s responses to stimulus *k* as *r_1_(k,n)* on the first repeat and *r_2_(k,n)* on the second repeat. These responses contain a stimulus component *s(k,n)* and some noise, where the stimulus component is defined as the trial-averaged response of neuron *n* to stimulus *k,* for an infinite number of repeated identical trials. We can thus write

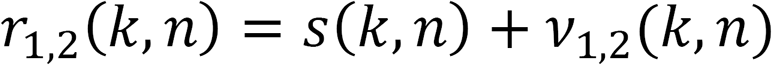

where *v*_1,2_ is the independent single trial variance. We compute a measure of stimulus tuning defined as the variance of *s(k, n)* over *k*, which we write *Var_k_*(*s*(*k, n*)). We can obtain this quantity by observing that

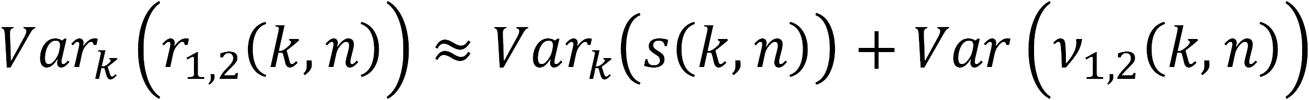

because the variance can be taken first over the random noise variable, and then over the stimulus dimension. We also have

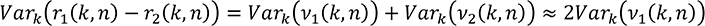

because the random variables *v*_1,2_ are independent of each other, have mean zero, and have the same variance. It follows that the fraction stimulus-related variance *Var_k_*(*s*(*k,n*)) can be computed by

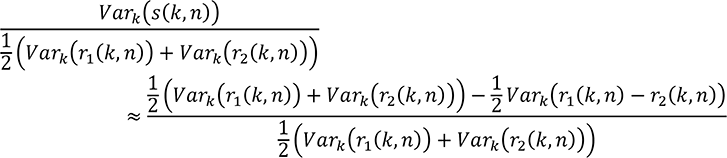

This latter quantity is approximately equal to the Pearson correlation *σ_stim_* of *r*_1_(*k,n*) and *r*_2_(*k,n*), allowing for the replacement of the arithmetic mean at the denominator with the geometric mean of the two variances that are in the limit equal to each other, and in practice very nearly so.

Note also that in practice we compute the rank correlation (Spearman) rather than the Pearson correlation, to avoid potential biases introduced by nonlinearities in the spike deconvolution process. For example, if the result of deconvolution to both repeats is transformed through the same nonlinearity, the Pearson correlation may be artificially increased, while the Spearman correlation remains the same.

### Convolutional neural networks

Of the SPIKEFINDER challenge winners, one team has so far made their code publicly available (https://github.com/PTRRupprecht/Spikefinder-Elephant/tree/master/elephant), so we used their CNN configuration, called “Elephant”, which has >100,000 free parameters.

## Acknowledgements

We thank Charu Reddy for assistance with surgeries and Michael Krumin for assistance with the two-photon microscopes. This work was supported by the Wellcome Trust (95668, 95669, 108726), and the Simons Foundation (325512). CS was funded by a four-year Gatsby Foundation PhD studentship.

